# Data-driven Discovery of Mathematical and Physical Relations in Oncology Data using Human-understandable Machine Learning

**DOI:** 10.1101/2021.08.13.456200

**Authors:** Daria Kurz, Carlos Salort Sánchez, Cristian Axenie

## Abstract

For decades, researchers have used the concepts of rate of change and differential equations to model and forecast neoplastic processes. This expressive mathematical apparatus brought significant insights in oncology by describing the unregulated proliferation and host interactions of cancer cells, as well as their response to treatments. Now, these theories have been given a new life and found new applications. With the advent of routine cancer genome sequencing and the resulting abundance of data, oncology now builds an “arsenal” of new modeling and analysis tools. Models describing the governing physical laws of tumor-host-drug interactions can be now challenged with biological data to make predictions about cancer progression. Our study joins the efforts of the mathematical and computational oncology community by introducing a novel machine learning system for data-driven discovery of mathematical and physical relations in oncology. The system utilizes computational mechanisms such as competition, cooperation, and adaptation in neural networks to simultaneously learn the statistics and the governing relations between multiple clinical data covariates. Targeting an easy adoption in clinical oncology, the solutions of our system reveal human-understandable properties and features hidden in the data. As our experiments demonstrate, our system can describe nonlinear conservation laws in cancer kinetics and growth curves, symmetries in tumor’s phenotypic staging transitions, the pre-operative spatial tumor distribution, and up to the nonlinear intracellular and extracellular pharmacokinetics of neoadjuvant therapies. The primary goal of our work is to enhance or improve the mechanistic understanding of cancer dynamics by exploiting heterogeneous clinical data. We demonstrate through multiple instantiations that our system is extracting an accurate human-understandable representation of the underlying dynamics of physical interactions central to typical oncology problems. Our results and evaluation demonstrate that using simple - yet powerful - computational mechanisms, such a machine learning system can support clinical decision making. To this end, our system is a representative tool of the field of mathematical and computational oncology and offers a bridge between the data, the modeler, the data scientist, and the practising clinician.

## 1 INTRODUCTION

The dynamics governing cancer initiation, development, and response to treatment are informed by quantitative measurements. These measurements carry details about the physics of the underlying processes such as, tumor growth, tumor-host cells encounters, and drug transport. Be it through mathematical modeling and patient-specific treatment trajectories — as in the excellent work of Werner et al. (2016) — through tumor’s mechano-pathology — systematically described by Nia et al. (2020a)— or through hybrid modeling frameworks of tumor development and treatment — identified by Chamseddine and Rejniak (2020)— capturing such processes from data can substantially improve predictions about cancer progression.

Machine learning algorithms are now leveraging automatic discovery of physics principles and governing mathematical relations for such improved predictions. Proof stands the proliferating body of such research — for representative results see the work of Raissi (2018), Schaeffer (2017), Long et al. (2018), and Champion et al. (2019)). However, the naive application of such algorithms is insufficient to infer physical laws underlying cancer progression. Simply positing a physical law or mathematical relation from data is useless without simultaneously proposing an accompanying ground truth to account for the inevitable mismatch between model and observations, as demonstrated in the work of de Silva et al. (2020).

Such a problem is even more important in clinical oncology where, in order to understand the links between the physics of cancer and signaling pathways in cancer biology, we need to describe the fundamental physical principles shared by most if not all tumors, as proposed in Nia et al. (2020b). Here, mathematical models of the physical mechanisms and corresponding tumor physical hallmarks complement the heterogeneity of the experimental observations. Such a constellation is typically validated through in vivo and in vitro model systems where the simultaneous identification of both the structure and parameters of the dynamical system describing tumor-host interactions is performed (White et al. (2019)).

Given the multi-dimensional nature of this system identification process, some concepts involved are non-intuitive and require deep and broad understanding of both the physical and biological aspects of cancer. To circumvent this, combining mechanistic modelling and machine learning is a promising approach with high potential for clinical translation. For instance, in a bottom-up approach, fusing cell-line tumor growth curve learning from heterogeneous data (i.e. caliper, imaging, microscopy) and unsupervised extraction of cytostatic pharmacokinetics, the study of Axenie and Kurz (2020a) introduced a novel pipeline for patient-tailored neoadjuvant therapy planning. In another relevant study, Benzekry (2020) used machine learning to extract model parameters from high-dimensional baseline data (demographic, clinical, pathological molecular) and employed mixed effects theory to combine it with mechanistic models based on longitudinal data (e.g. tumor size measurements, pharmacokinetics, seric biomarkers, and circulating DNA) for treatment individualization.

Yet, despite the recent advances in mathematical and computational oncology, there are only a few systems trying to offer a human understandable solution, or the steps to reach it — the most relevant are the studies of Jansen et al. (2020) and Lamy et al. (2019). But, such systems lack a rigorous and accessible description of the physical cancer traits assisting their clinical predictions. Our study advocates the improvement of mechanistic modelling with the help of machine learning. Our thesis goes beyond measurements-informed bio-physical processes models, as described in Cristini et al. (2017), and towards human-understandable personalized disease evolution and therapy profiles learnt from data, as foreseen in Kondylakis et al. (2020).

### 1.1 Study Focus

The purpose of this study is to introduce a system (and a framework) capable of learning human-understandable mathematical and physical relations from heterogeneous oncology data for patient-centered clinical decision support. To demonstrate the versatility of the system, we introduce multiple of its instantiations, in an end-to-end fashion (i.e. from cancer initiation to treatment outcome) for predictions based on available clinical datasets^1^:

- **learning initiation patterns of pre-invasive breast cancer** (i.e. Ductal Carcinoma In-Situ, DCIS) from histopathology and morphology data available from the studies of Rodallec et al. (2019); Volk et al. (2011); Tan et al. (2015) and Mastri et al. (2019);
- **learning unperturbed tumor growth curves within and between cancer types** (i.e. breast, lung, leukemia) from imaging, microscopy, and caliper data available from the studies of Benzekry et al. (2019) and Simpson-Herren and Lloyd (1970);
- **extracting tumor phenotypic stage transitions** from three cell lines of breast cancer using imaging, immunohistochemistry, and histopathology data available from the studies of Rodallec et al. (2019); Volk et al. (2011); Tan et al. (2015) and Edgerton et al. (2011);
- **simultaneously extracting the drug-perturbed tumor growth and drug pharmacokinetics** for neoadjuvant/adjuvant therapy sequencing using data available from the studies of Kuh et al. (2000); Volk et al. (2011) and Chen et al. (2014);
- **predicting tumor growth/recession** (i.e. estimating tumor volume after each chemotherapy cycle **under various chemotherapy regimens** administered to breast cancer patients, using real-world patient data available from the study of Yee et al. (2020) as well as cell lines studies from Rodallec et al. (2019); Volk et al. (2011); Tan et al. (2015) and Mastri et al. (2019).

In each of the instantiations, we use the same computational substrate (i.e. no specific task parametrization) and compare the performance of our system against state-of-the-art systems capable of extracting governing equations from heterogeneous oncology data from of Cook et al. (2010), Mandal and Cichocki (2013), Weber and Wermter (2007), and Champion et al. (2019), respectively. The analysis focuses on: 1) the accuracy of the systems in the learnt mathematical and physical relations among various covariates, 2) the ability to embed more data and mechanistic models, and 3) the ability to provide a human-understandable solution and the processing steps to obtain that solution.

### 1.2 Study Motivation

In clinical practice, patient tumors are typically described across multiple dimensions: from 1) high-dimensional heterogeneous data (e.g. demographic, clinical, pathological, molecular), and 2) longitudinal data (e.g. tumor size measurements, pharmacokinetics, immune screening, biomarkers), to 3) time-to-event data (e.g. progression-free or overall survival analysis), and, in the last years, 4) genetic sequencing that determine the genetic mutations driving their cancer. With this information, the clinical oncologist may tailor treatment to the patient’s specific cancer.

But, despite the variety of such rich patient data available, tumor growth data, describing the dynamics of cancer development, from initiation to metastasis has some peculiarities. These features motivated the study and the approach proposed by our system. To summarize, tumor growth data:

- is typically **small**, with only a few data points measured, typically, at days-level resolution (Roland et al. (2009));
- is **unevenly sampled**, with irregular spacing among tumor size/volume observations (Volk et al. (2011));
- has **high variability** between and within tumor types (Benzekry et al. (2014)) and type of treatment (Gaddy et al. (2017)).
- is **heterogeneous** and sometimes **expensive or difficult to obtain** (e.g. bio-markers, fMRI (Abler et al. (2019)), fluorescence imaging (Rodallec et al. (2019)), flow cytometry, or calipers (Benzekry et al. (2019)).
- **determines cancer treatment planning**, for instance, adjuvant vs. neoadjuvant chemotherapy, (Sarapata and de Pillis (2014)).

Using unsupervised learning, our system seeks to overcome these limitations and provide a human-understandable representation of the mathematical and physical relations describing tumor growth, its phenotype, and, finally, its interaction with chemotherapeutic drugs. The system exploits the temporal evolution of the processes describing growth data along with their distribution in order to reach superior accuracy and versatility on various clinical in-vitro tumor datasets.

## 2 MATERIALS AND METHODS

In the current section, we introduce our system through the lens of practical examples of discovering mathematical and physical relations describing tumor-host-drug dynamics. We begin by introducing the basic computational framework as well as the various configurations in which the system can be used. The second part is dedicated to introducing relevant state-of-the-art approaches used in our comparative experimental evaluation.

### 2.1 System Basics

Physical interactions of cancer cells with their environment (e.g. local tissue, immune cells, drugs) determine the physical characteristics of tumors through distinct and interconnected mechanisms. For instance, cellular proliferation and its inherent abnormal growth patterns lead to increased solid stress (Nia et al. (2016)). Subsequently, cell contraction and cellular matrix deposition modify the architecture of the surrounding tissue which can additionally react to drugs (Griffon-Etienne et al. (1999)) modulating the stiffness (Rouvière et al. (2017)) and interstitial fluid pressure (Nathanson and Nelson (1994)). But such physical characteristics also interact among each other initiating complex dynamics, as demonstrated in Nia et al. (2020a).

Our system can capture such complex dynamics through a network-based paradigm for modelling, computation, and prediction. It can extract the mathematical description of the interactions exhibited by multiple entities (e.g. tumor, host cells, cytostatic drugs) for producing informed predictions. For guiding the reader, we present a simple, biologically grounded example in Figure 1. In this example, our system learns simultaneously the power-law tumor growth under immune escape (Benzekry et al. (2014)) and the nonlinear potentiation-inhibition model of Natural Killer Cells (NK)-Tumor interactions (Ben-Shmuel et al. (2020)), while exhibiting the known overlapping Cytotoxic T Lymphocytes (CTL)-Natural Killer Cells (NK) mutual linear regulation pattern (Uzhachenko and Shanker (2019)). As shown in Figure 1, our system offers the means to learn the mathematical relations governing the physical tumor-immune interactions, without supervision, from available clinical data (see Figure 1 - input data relations and learned and decoded relations). Furthermore, the system can infer unavailable (i.e. expensive to measure) physical quantities (i.e. after learning/training) in order to make predictions on the effects of modifying the pattern of interactions among the tumor and the immune system. For instance by not feeding the system with the innate immune response (i.e. the NK cells dynamics), the system infers, based on the CTL-NK interaction pattern and the tumor growth pattern, a plausible tumor-NK mathematical relation in agreement with observations (see Figure 1 - left panel, squared root non-linearity).

**Figure 1.**
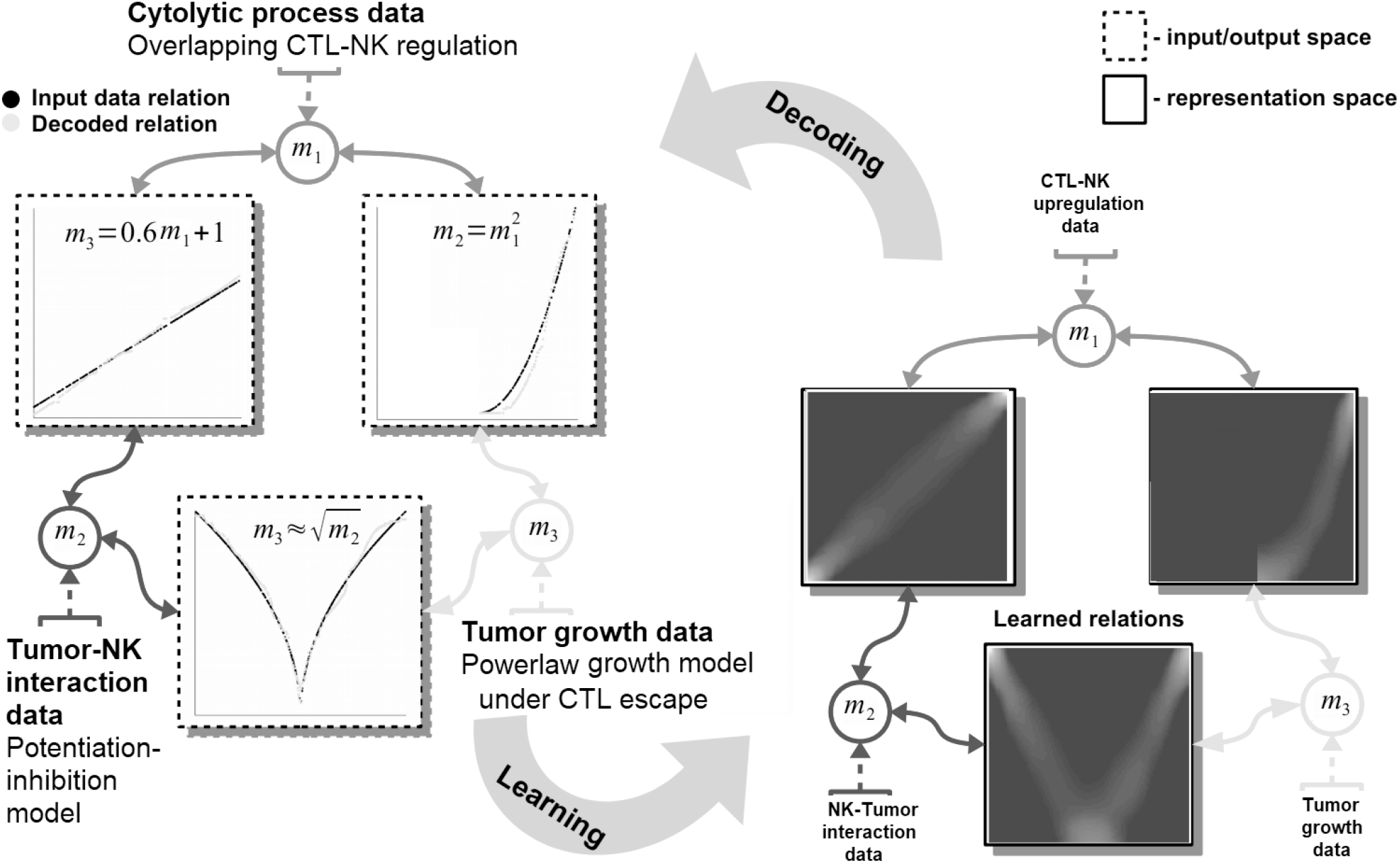
Basic functionality of the proposed machine learning system. Data is fed in the system through the representations maps, *m*_*i*_ which encode each quantity in a distributed (array-like) representation. The system dynamics brings all available quantities into agreement and learns the underlying mathematical relations among them (see representation space - right panel). The relations resemble the mathematical model of the interactions: power-law tumor growth under immune escape, nonlinear potentiation-inhibition tumor-immune interaction, and linear regulation pattern among immune system cells. The learnt mathematical relations are then compared with the mechanistic model output, considered ground truth (see input/output space - left panel).

Basically, our system acts as constraint satisfaction network converging to a global consensus given local (i.e. the impact of the measured data) and global dynamics of the physics governing the interactions (see the clear patterns depicting the mathematical models of interaction in Figure 1). The networked structure allows the system to easily interconnect multiple data quantities measuring different biological components (Markowetz and Troyanskaya (2007)) or a different granularity of representation of the underlying interaction physics (Cornish and Markowetz (2014)).

### 2.2 Computational Substrate

The core element of our study is an unsupervised machine learning system based on Self-Organizing Maps (SOM) Kohonen (1982) and Hebbian Learning (HL) Chen et al. (2008). The two components are used in concert to represent and extract the underlying relations among correlated data. In order to introduce the computational steps followed by our system, we provide a simple example in Figure 2. Here, we feed the system with data from a cubic growth law (3^*rd*^ power-law) describing the effect of drug dose density over 150 weeks of adjuvant chemotherapy in breast cancer (data from Comen et al. (2016)). The two data sources (i.e. the cancer cell number and the irregular measurement index over the weeks) follow a cubic dependency (cmp. Figure 2a). Before being presented the data, our system has no prior information about the data distribution and its generating process (or model). The system learns the underlying (i.e. hidden) mathematical relation directly from the pairs of input data without supervision. The input SOMs (i.e. 1D lattice networks with *N* neurons) extract the probability distribution of the incoming data, depicted in Figure 2a, and encode samples in a distributed activity pattern, as shown in Figure 2b. This activity pattern is generated such that the closest preferred value of a SOM neuron to the input will be strongly activated and will decay, proportional with distance, for neighbouring units. This process is fundamentally benefiting from the quantization capability of SOM. Additionally, this process is extended with a dimension corresponding to the latent representation of network resource allocation (i.e. number of neurons allocated to represent the input data space). After learning, the SOMs specialise to represent a certain (preferred) value in the input data space and learns its probability distribution, by updating its tuning curves shape. Practically, given an input value *s*^*p*^(*k*) from one timeseries at time step *k*, the network follows the processing stages in Figure 3. For each *i*-th neuron in the *p*-th input SOM, with preferred value 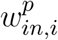 and tuning curve size 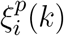, the generated neural activation is given by

**Figure 2.**
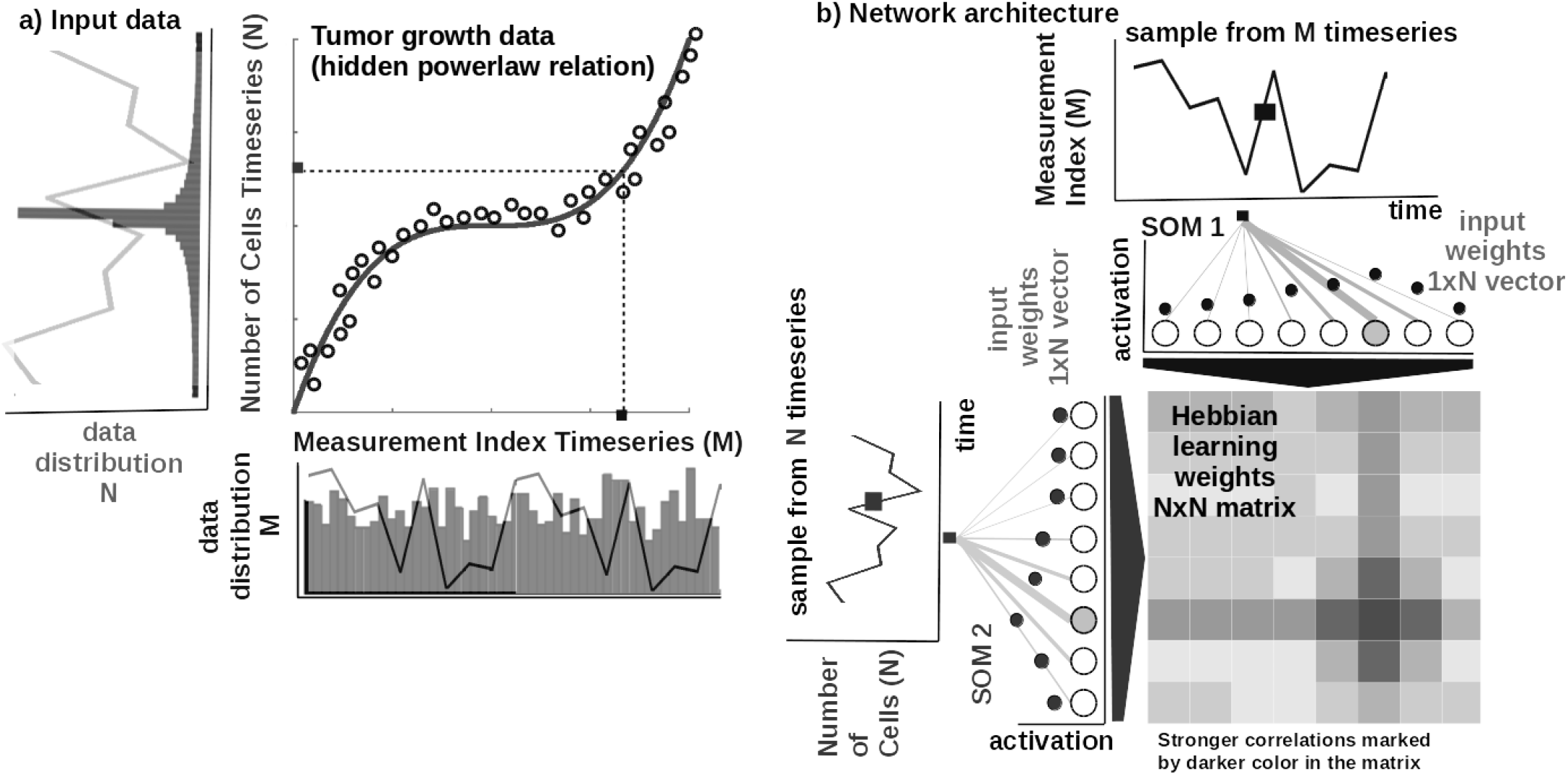
Basic functionality of the system. a) Tumor growth data following a non-linear mathematical relation and its distribution - relation is hidden in the timeseries (i.e. number of cells vs. measurement index). Data from Comen et al. (2016). b) Basic architecture of our system: 1-dimensional (array) SOM networks with *N* neurons encoding the timeseries (i.e. number of cells vs. measurement index), and a *N × N* Hebbian connection matrix (coupling the two SOMs) that will encode the mathematical relation after training.

**Figure 3.**
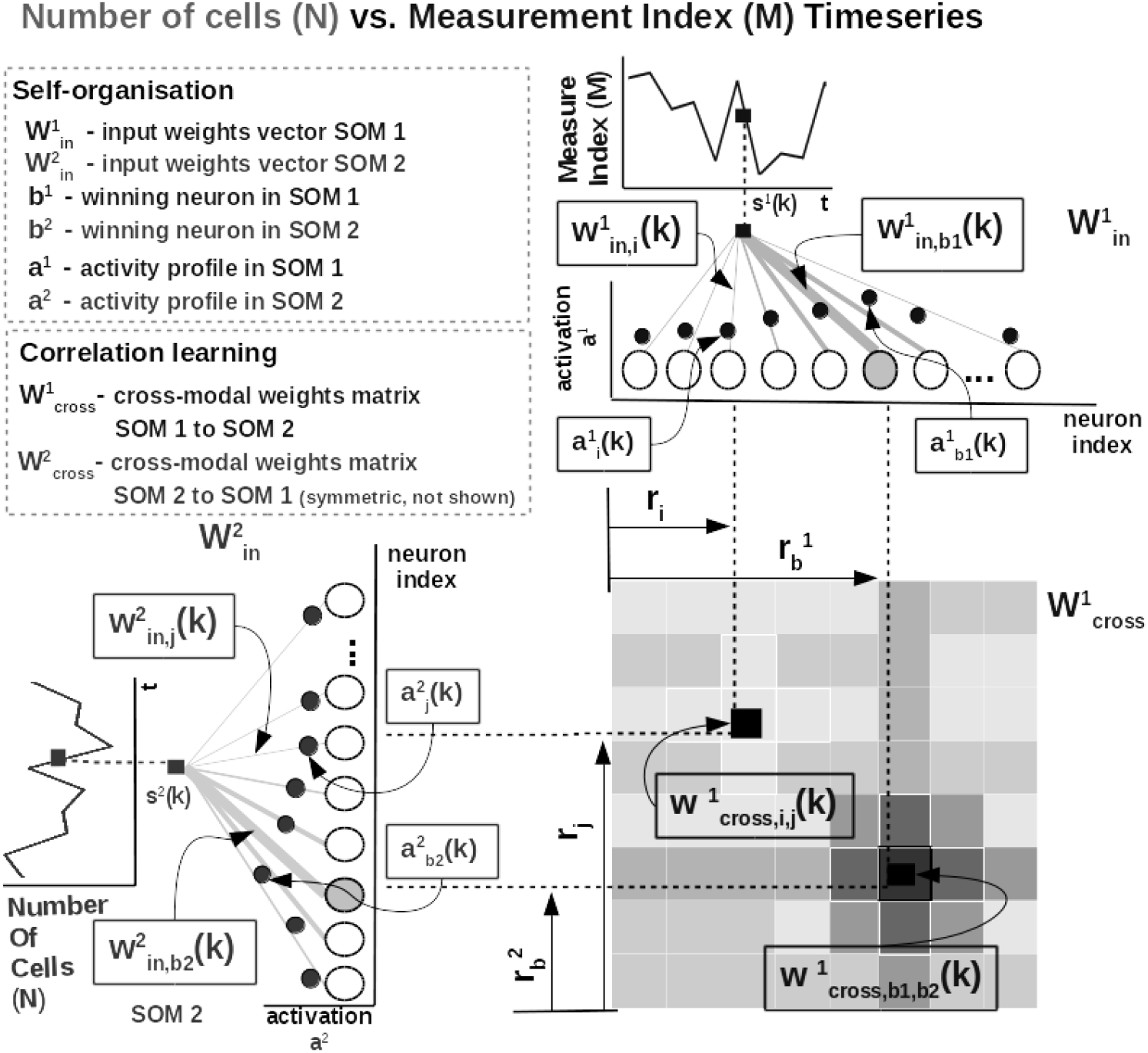
Detailed computational steps of our system, instantiated for tumor growth learning given the observed number of cells and the measurement index data from Comen et al. (2016).

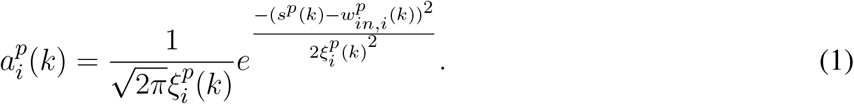

The most active (i.e. competition winning) neuron of the *p*-th population, *b*^*p*^(*k*), is the one which has the highest activation given the timeseries data point at time *k*

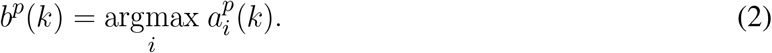

The competition for highest activation (in representing the input) in the SOM is followed by a cooperation process that captures the input space distribution. More precisely, given the winning neuron, *b*^*p*^(*k*), the cooperation kernel,

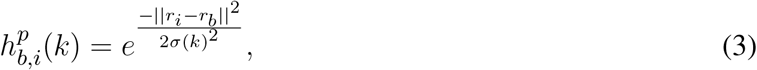

allows neighbouring neurons in the network (i.e. found at position *r*_*i*_ in the network) to precisely represent the input data point given their location in the neighbourhood *σ*(*k*) of the winning neuron. The topological neighbourhood width *σ*(*k*) decays in time, to avoid artefacts (e.g. twists) in the SOM. The kernel in Equation 3, is chosen such that adjacent neurons in the network specialise on adjacent areas in the input space, by “pulling” the input weights (i.e. preferred values) of the neurons closer to the input data point,

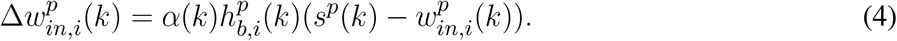

This process updates the tuning curves width 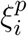 given the spatial location of the neuron in the network, the distance to the input data point, the cooperation kernel size, and a decaying learning rate *α*(*k*),

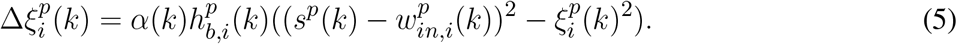

To illustrate these mechanisms, we consider the learned tuning curves shapes for 5 neurons in the input SOMs (i.e. neurons 1, 6, 13, 40, 45) encoding the breast cancer cubic tumor growth law, depicted in Figure 4. We observe that higher input probability distributions are represented by dense and sharp tuning curves (e.g. neuron 1, 6, 13 in SOM1), whereas lower or uniform probability distributions are represented by more sparse and wide tuning curves (e.g. neuron 40, 45 in SOM1). This way, the system optimally allocates neurons such that a higher amount of neurons represent areas in the input space, which need a finer resolution; and a lower amount for more coarsely represented input space areas. Neurons in the two SOMs are then linked by a fully (all-to-all) connected matrix of synaptic connections, where the weights are computed using Hebbian learning. The connections between uncorrelated (or weakly correlated) neurons in each SOM (i.e. *w*_*cross*_) are suppressed (i.e. darker color) while correlated neurons connections are enhanced (i.e. brighter color), as depicted in Figure 3. Each connection weight 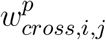 between neurons *i, j* in the input SOMs are updated with a Hebbian learning rule as follows:

**Figure 4.**
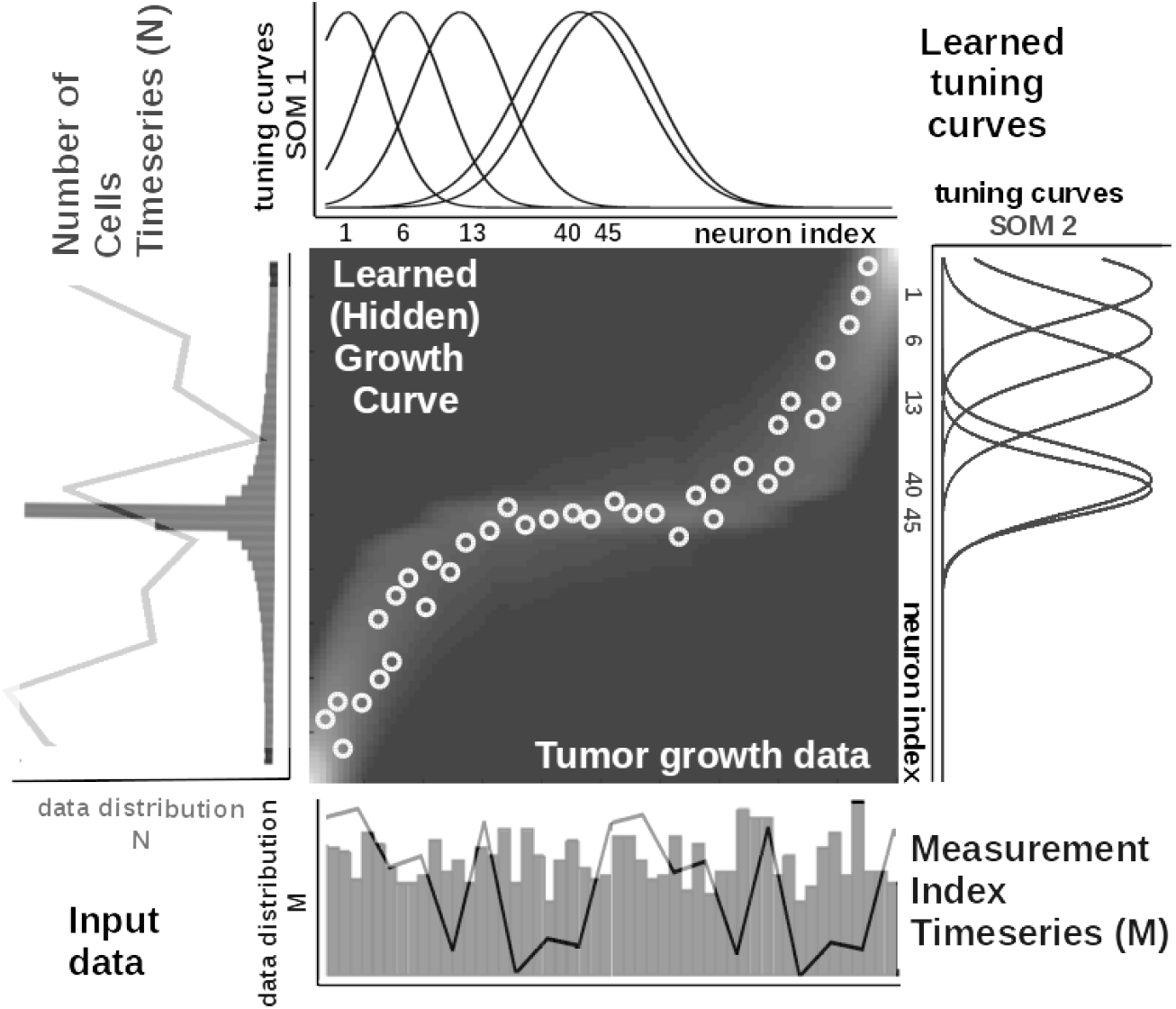
Extracted mathematical relation describing the growth law and data statistics for the experimental observations in Figure 2a depicting a cubic breast cancer tumor growth law among number of cells and irregular measurement over 150 weeks from Comen et al. (2016). Raw data timeseries is overlaid on the data distribution and corresponding model encoding tuning curves shapes.

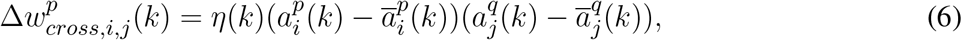

where

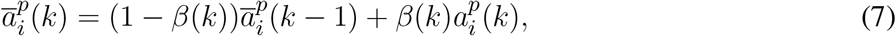

is an exponential decay (i.e. momentum) and *η*(*k*), *β*(*k*) are monotonic (inverse-time) decaying functions. Hebbian learning ensures a weight increase for correlated activation patterns and a weight decrease for anti-correlated activation patterns. The Hebbian weight matrix encodes the co-activation patterns between the input SOMs, as shown in Figure 2b, and, eventually, the learned mathematical relation given the data, as shown in Figure 4. Such a representation, as shown in Figure 4, demonstrates the human-understandable output of our system which employs powerful, yet simple and transparent, processing principles, as depicted in Figure 3.

Input SOM self-organisation and Hebbian correlation learning operate at the same time in order to refine both the input data representation and the extracted mathematical relation. This is visible in the encoding and the decoding functions where the input activations *a* are projected through the input weights *w*_*in*_ (Equation 1) to the Hebbian matrix and then decoded through the *w*_*cross*_ correlation weights. (Equation 8).

In order to recover the real-world value from the network, we use a decoding mechanism based on (self-learnt) bounds of the input data space. The input data space bounds are obtained as minimum and maximum of a cost function of the distance between the current preferred value of the winning neuron (i.e. the value in the input which is closest (in Euclidian space) to the weight vector of the neuron) and the input data point in the SOM (i.e. using Brent’s optimization Brent (2013)). Depending on the position of the winning neuron in the SOM, the decoded / recovered value *y*(*t*) from the SOM neurons weights is computed as:

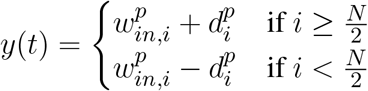

where, 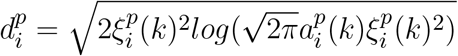 for the winning neuron with index *i* in the SOM, a preferred value 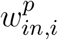 and 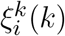 tuning curve size and 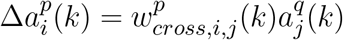. The activation 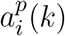 is computed by projecting one data point through SOM *q* and subsequently through the Hebbian matrix to compute the paired activity (i.e. at the other SOM *p*, Equation 8) describing the other data quantity.

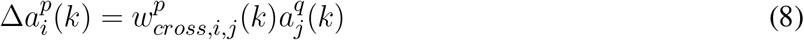

where 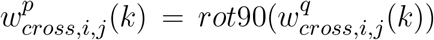 and *rot*90 is a clockwise rotation. The decoding step, is a fundamental aspect contributing to the human-understandable output of our system. This demonstrates that simple operations, such as competition and cooperation in neural networks can exploit the statistics of clinical data and provide a human-understandable representation of the governing mathematical relations behind tumor growth processes.

### 2.3 Comparable Systems

In this section, we briefly introduce five state-of-the-art approaches that we comparatively evaluated against our system. Ranging from statistical methods, to machine learning, and up to deep learning, the selected systems were designed to extract governing equations from data.

#### Cook et al

The system of Cook et al. (2010) uses a combination of simple computational mechanisms, like Winner-Take-All (WTA) circuits, Hebbian learning, and homeostatic activity regulation, to extract mathematical relations among different data sources. Real-world values presented to the network are encoded in population code representations. This approach is similar to our approach in terms of the sparse representation used to encode data. The difference resides in the fact that in our model the input population (i.e. SOM network) connectivity is learned. Using this capability, our model is capable of learning the input data bounds and distribution directly from the input data, without any prior information or fixed connectivity. Furthermore, in this system, the dynamics between each population encoded input is performed through plastic Hebbian connections. Starting from a random connectivity pattern, the matrix finally encoded the functional relation between the variables which it connects. The Hebbian linkage used between populations is the correlation detection mechanism used also in our model, although in our formulation we adjusted the learning rule to accommodate both the increase and decrease of the connection weights.

#### Weber et al

Using a different neurally inspired substrate, the system of Weber and Wermter (2007) combines competition and cooperation in a self-organizing network of processing units to extract coordinate transformations. More precisely, the model uses simple, biologically motivated operations, in which co-activated units from population coded representations self-organize after learning a topological map. This basically assumes solving the reference frame transformation between the inputs (mapping function). Similar to our model the proposed approach extends the SOM network by using sigma-pi units (i.e. weighted sum of products). The connection weight between this type of processing units implements a logical AND relation.The algorithm produces invariant representations and a topographic map representation.

#### Mandal et al

Going away from biological inspiration, the system of Mandal and Cichocki (2013) used a type of nonlinear canonical correlation analysis (CCA), namely alpha-beta divergence correlation analysis (ABCA). The ABCA system extracts relations between sets of multidimensional random variables. The core idea of the system is to first determine linear combinations of two random variables (called canonical variables/variants) such that the correlation between the canonical variables is the highest amongst all such linear combinations. As traditional CCA is only able to extract linear relations between two sets of multi-dimensional random variable, the proposed model comes as an extension to extract nonlinear relations, with the requirement that relations are expressed as smooth functions and can have a moderate amount of additive random noise on the mapping. The model employs a probabilistic method based on nonlinear correlation analysis using a more flexible metric (i.e. divergence / distance) than typical canonical correlation analysis.

#### Champion et al

As Deep Learning (DL) is becoming a routine tool for data discovery, as shown in the recent work of Champion et al. (2019); Raissi (2018); Schaeffer (2017), and de Silva et al. (2020), we also consider a DL system (inspired from Champion et al. (2019)) and evaluate it along the other methods. To apply this prediction method to tumor growth, we need to formulate the setup as a timeseries prediction problem. At any given point, we have the dates and values of previous observations. Using these two features, we can implement DL architectures that predict the size of the tumor at a future step. Recurrent Neural Networks (RNN) are the archetypal DL architectures for timeseries prediction. The principal characteristic of RNN, compared to simpler DL architectures, is that they iterate over the values that have been observed, obtaining valuable information from it, like the rate at which the objective variable grows, and use that information to improve prediction accuracy. The main drawback of using DL in the medical field is the need of DL models to be presented with large amounts of data. We address this problem by augmenting the data. We use Support Vector Machines (SVM) for augmenting data, to obtain expected tumor development with normal noise generates realistic measurements. This approach presents models the expected average development of a tumor.

## 3 EXPERIMENTAL SETUP AND RESULTS

In order to evaluate our data-driven approach to learn mathematical and physical relations from heterogeneous oncology data, we introduce the five instantiations and their data briefly introduced in the Study Focus section.

### 3.1 Datasets

In our experiments, we used publicly available tumor growth, pharmacokinetics, and chemotherapy regimens datasets (see Table 1), with in-vitro or in-vivo clinical tumor volume measurements, for breast cancer (datasets 1, 2, 5, 6, 7) and other cancers (e.g. lung, leukemia - datasets 3 and 4, respectively). This choice is to probe and demonstrate transfer capabilities of the system to tumor growth patterns induced by different cancer types.

**Table 1.** Description of the datasets used in the experiments. 1 - dataset from the study of Rodallec et al. (2019) 2 - dataset from the study of Volk et al. (2011) 3 - dataset from the study of Benzekry et al. (2019) 4 - dataset from the study of Simpson-Herren and Lloyd (1970) 5 - dataset from the study of Tan et al. (2015) 6 - dataset from the study of Mastri et al. (2019) 7 - dataset from the study of Yee et al. (2020)

| Experimental dataset setup |  |  |  |  |
| --- | --- | --- | --- | --- |
| Dataset | Cancer Type | Data Type | Data Points | Data Freq. |
| 1 | Breast<sup>1</sup>(MDA-MB-231 cell line) | Fluorescence imaging | 7 | 2x/week |
| 2 | Breast<sup>2</sup>(MDA-MB-435 cell line) | Digital Caliper | 14 | 2x/week |
| 3 | Lung<sup>3</sup> | Caliper | 10 | 7x/week |
| 4 | Leukemia<sup>4</sup> | Microscopy | 23 | 7x/week |
| 5 | Breast<sup>5</sup>(MCF-7 cell line) | Microscopic imaging | 8 | 1x/week |
| 6 | Breast<sup>6</sup>(LM2-4LUC+ cell line) | Digital Caliper | 10 | 3x/week |
| 7 | Breast<sup>7</sup>(stage 2/3 cancers) | fMRI | 5 | 1x/week |

For the pharmacokinetics experiments (i.e. mainly focused on taxanes family for experiments on MCF-7 breast cancer cell line from Tan et al. (2015)), we used the data from Kuh et al. (2000) describing intracellular and extracellular concentrations of Paclitaxel during uptake. The datasets and the code for all the systems used in our evaluation are available on GitLab^2^.

### 3.2 Procedures

#### Our system

In all of our experiments, data depicting tumor growth, pharmacokinetics, and chemotherapy regimens is fed to our system which encodes each timeseries in the SOMs and learns the underlying relations in the Hebbian matrix. The SOMs are responsible of bringing the timeseries in the same latent representation space where they can interact (i.e. through their internal correlation). Throughout the experiments, each of the SOM has *N* = 100 neurons, the Hebbian connection matrix has size *N × N* and parametrization is done as: *α* = [0.01, 0.1] decaying, 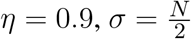 decaying following an inverse time law.

#### Cook et al

For the neural network system proposed by Cook et al. (2010), in all our experiments, we used neural populations with 200 neurons each, a 0.001 WTA settling threshold, 0.005 scaling factor in homeostatic activity regulation, 0.4 amplitude target for homeostatic activity regulation, and 250 training epochs. More details and the reference codebase is available on GitLab.

#### Weber et al

For the neural network system proposed by Weber and Wermter (2007), in all our experiments, we used a network with 15 neurons, 0.001 learning rate, 200000 training epochs, and unit normalization factor. The fully parametrized codebase is available, along the other systems reference implementations, on GitLab.

#### Mandal et al

For the CCA-based system proposed by Mandal and Cichocki (2013), in all our comparative experiments, we used a sample size of 100, replication factor 10, 0.5 divergence factor, 1000 variable permutations, and 1.06 bandwidth for Gaussian kernel density estimate. The full codebase is provided, along the other systems reference implementations, on GitLab.

#### Champion et al

For our DL implementation we use the system of Champion et al. (2019) as a reference. We then modified the structure to accommodate the peculiarities of the clinical data. In all the experiments, the DL system contained hidden layers of size 128 neurons, trained for 100 epochs, with a mini-batch size of 1, and 50% augmentation percentage. The full codebase is provided, along the other systems reference implementations, on GitLab. Another important implementation aspect is that we use a combination of SVM and DL approaches. While SVM can work with a limited amount of data, DL models tend to perform worse when big data is not available. Therefore, we test multiple approaches to artificially augment the training data:

- **DL with no augmentation, DL**. We train the model directly from the data without further transformations.
- **DL with SVM augmentation, DL + SVM**. We used the SVM model trained beforehand to enhance the data. We set a number of observations that we want to enhance, and generate random timestamps we use for prediction using SVM. Then we add those artificial values as new observations for training.
- **DL with SVM augmentation and random noise, DL + SVM + noise**. We follow the same process as in SVM augmentation, but before adding the predictions to the training pool we add normal noise.

For the SVM we use one input feature, the days passed, and one output feature, the size of the tumor. For DL, we use Gated Recurrent Units (GRU) Chung et al. (2014) as building blocks to design a structure inspired by the work of Champion et al. (2019). The architecture consists on one GRU layer, one ReLU activation, a fully connected layer and another ReLU activation. We designed a simple architecture to better suit the model to the scarce availability of data inspired by the study of Berg and Nyström (2019). As the DL model is a recurrent model, our input data consists of all data available from a certain patient up to a point. Both models normalize the data (both days and tumor size) by dividing by the maximum value observed. We run a 4 fold cross validation for each dataset (except dataset 0, that has only 2 samples, therefore we run a 2 fold cross validation). We present the average results over the cross validation. The complete parametrization and implementation is available on GitLab.

### 3.3 Results

As previously mentioned, we evaluate the systems on a series of instantiations depicting various decision support tasks relevant for clinical use. All of the five models were evaluated through multiple metrics (see Table 2) on each of the four cell line datasets. In order to evaluate the distribution of the measurement error as a function of the measured volumes of the tumors, the work of Benzekry et al. (2014) recommended the following model for the standard deviation of the error *σ*_*i*_ at each measurement time point *i*,

**Table 2.**
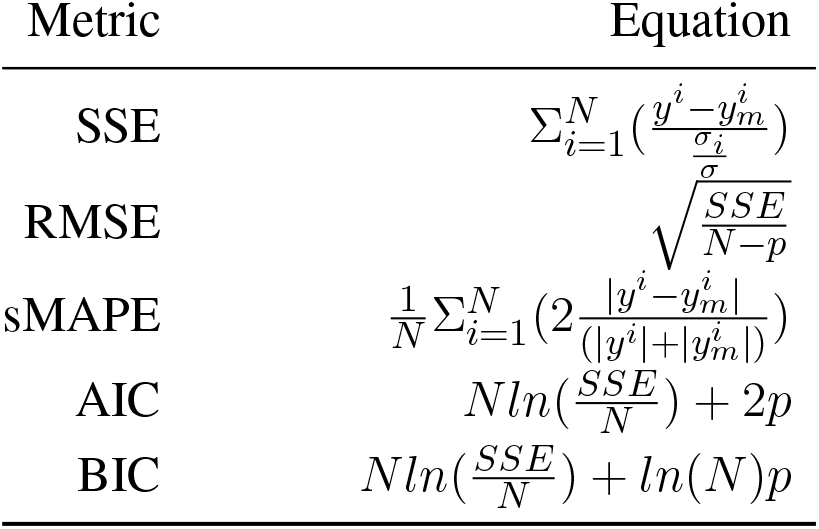
Evaluation metrics for data-driven relation learning systems. We consider: *N* - number of measurements, *σ* - standard deviation of data, *p* - number of parameters of the model.

**Table 3.**
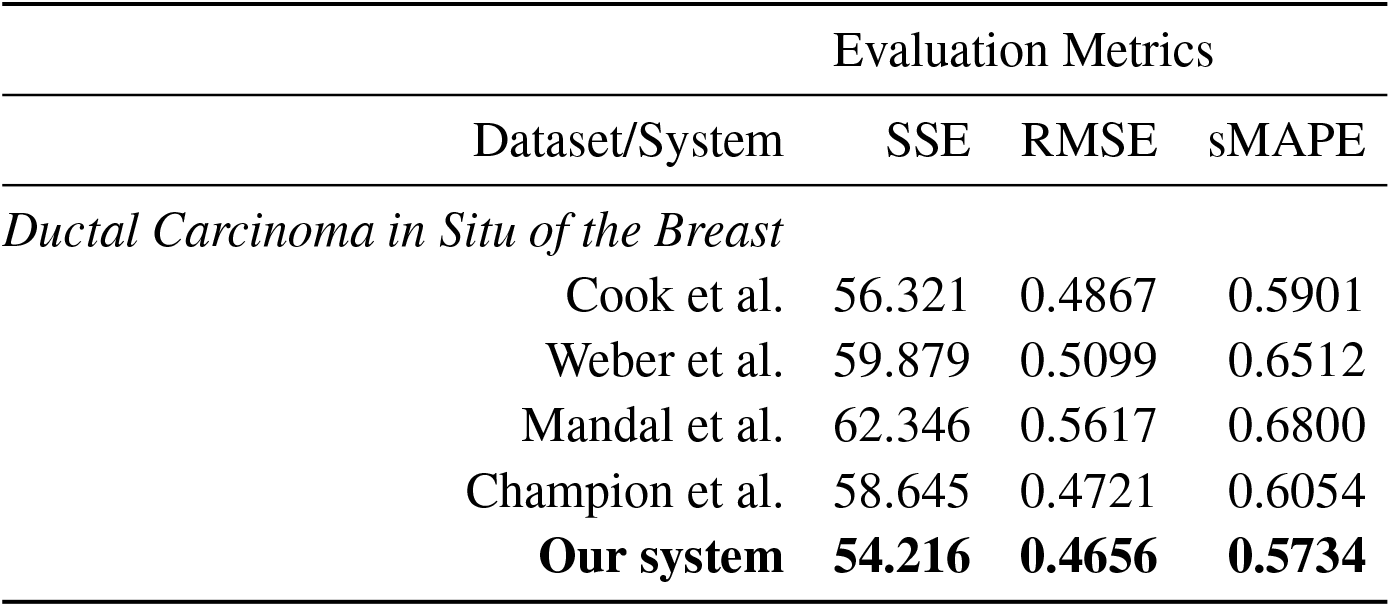
Evaluation of the data-driven relation learning systems.

|  |  | Evaluation Metrics |  |  |
| --- | --- | --- | --- | --- |
|  | Dataset/System | SSE | RMSE | sMAPE |
| *Ductal Carcinoma in Situ of the Breast* |  |  |  |  |
|  | Cook et al. | 56.321 | 0.4867 | 0.5901 |
|  | Weber et al. | 59.879 | 0.5099 | 0.6512 |
|  | Mandal et al. | 62.346 | 0.5617 | 0.6800 |
|  | Champion et al. | 58.645 | 0.4721 | 0.6054 |
|  | **Our system** | **54.216** | **0.4656** | **0.5734** |

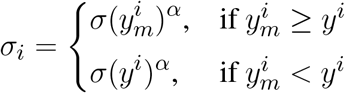

This model shows that when overestimating (*y*_*m*_ *≥ y*), the measurement error *α* is sub-proportional and, when underestimating (*y*_*m*_ *< y*), the obtained error is the same as the measured data points. In our experiments, we consider *α* = 0.84 and *σ* = 0.21, as a good trade-off of error penalty and enhancement. We use this measurement error formulation to calculate the typical performance indices (i.e. Sum of Squared Errors (SSE), Root Mean Squared Error (RMSE), Symmetric Mean Absolute Percentage Error (sMAPE)) and goodness-of-fit and parsimony (i.e. Akaike Information Criterion (AIC) and Bayesian Information Criterion (BIC)), as shown in Table 2.

#### 3.3.1 Learning growth patterns of pre-invasive breast cancer

Analysing tumor infiltration patterns, clinicians can evaluate the evolution of neoplastic processes, for instance from Ductal Carcinoma In Situ (DCIS) to breast cancer. Such an analysis can provide very important benefits, in early detection, in order to: (1) increase patient survival, (2) decrease the likelihood for multiple surgeries, and (3) determine the choice of adjuvant vs. neoadjuvant chemotherapy. For a full analysis and in-depth discussion of our system’s capabilities for such a task please refer to Axenie and Kurz (2020b). For this task, we assessed the capability of the evaluated systems to learn the dependency between histopathological and morphological data. We fed the systems with DCIS data from Edgerton et al. (2011), namely timeseries of nutrient diffusion penetration length within the breast tissue (*L*), ratio of cell apoptosis to proliferation rates (*A*), and radius of the breast tumor (*R*). The study of Edgerton et al. (2011) postulated that the value of *R* depends upon *A* and *L* following a “master equation” 9

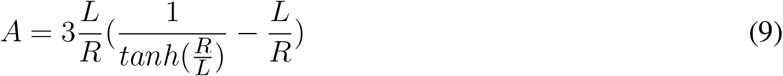

whose predictions are consistent with nearly 80% of in-situ tumors identified by mammographic screenings. For this initial evaluation of the data-driven mathematical relations learning systems, we consider three typical performance metrics (i.e. SSE, RMSE, and sMAPE, respectively) against the experimental data (i.e. ground truth and Equation 9):

As one can see, our system overcomes the other approaches on predicting the nonlinear dependency between radius of the breast tumor (*R*) given the nutrient diffusion penetration length within the breast tissue (*L*) and ratio of cell apoptosis to proliferation rates (*A*) from real in-vivo histopathological and morphological data.

#### 3.3.2 Learning unperturbed tumor growth curves within and between cancer types

In the second task, we evaluated the systems on learning unperturbed (i.e. growth without treatment) tumor growth curves. The choice of different cancer types (i.e. two breast cell lines, lung, and leukemia) is to probe and demonstrate between and within tumor type prediction versatility.

Our system provides overall better accuracy between and within tumor type growth curve prediction, as shown in Table 4 and the summary statistics (depicted in Figure 5). The superior performance is given by the fact that our system can overcome the other approaches when facing incomplete biological descriptions, the diversity of tumor types, and the small size of the data. Interested readers can refer to Axenie and Kurz (2021) for a deeper performance analysis of our system.

**Table 4.**
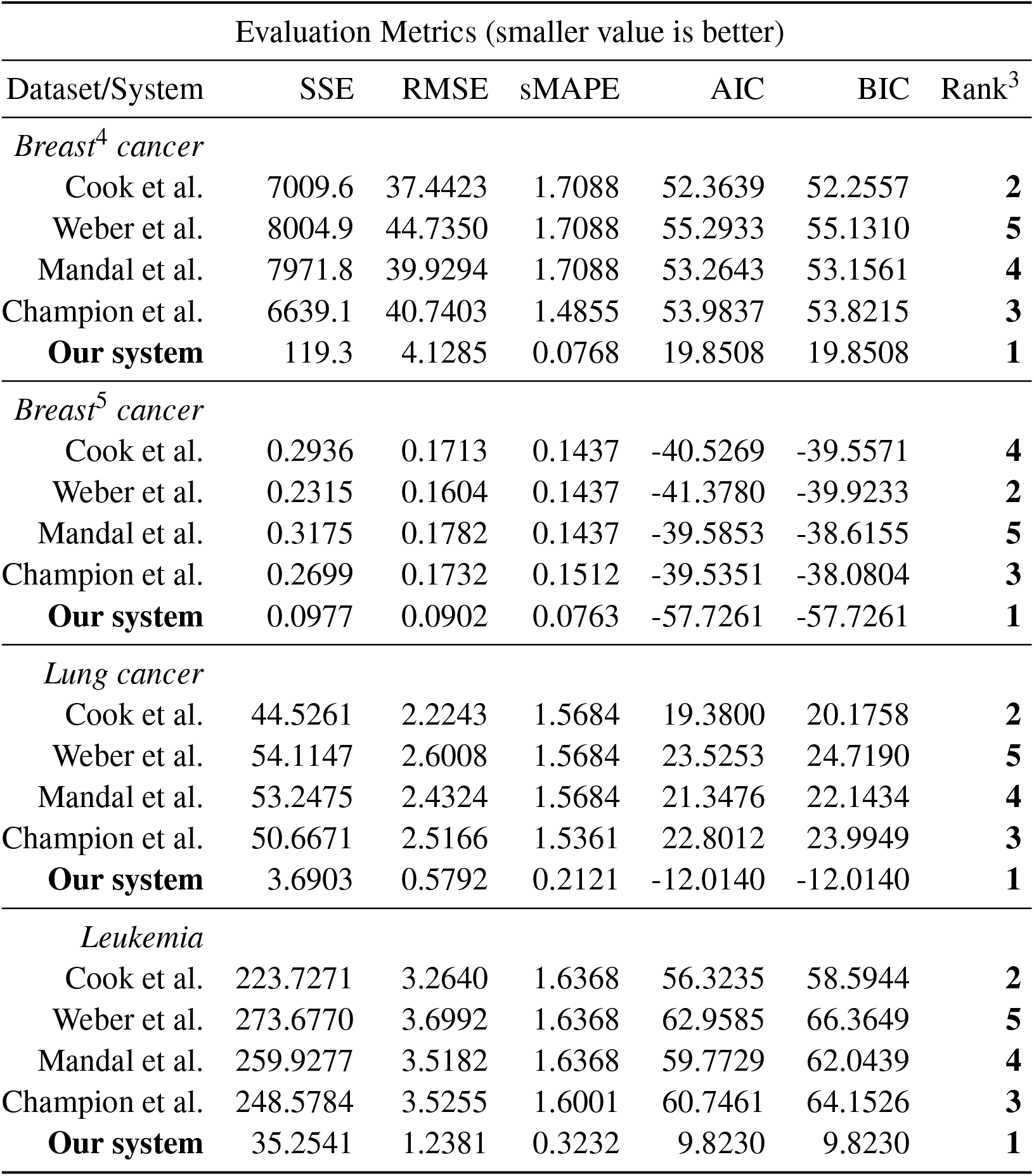
Evaluation of the data-driven relation learning systems on tumor growth curves extraction.

**Figure 5.**
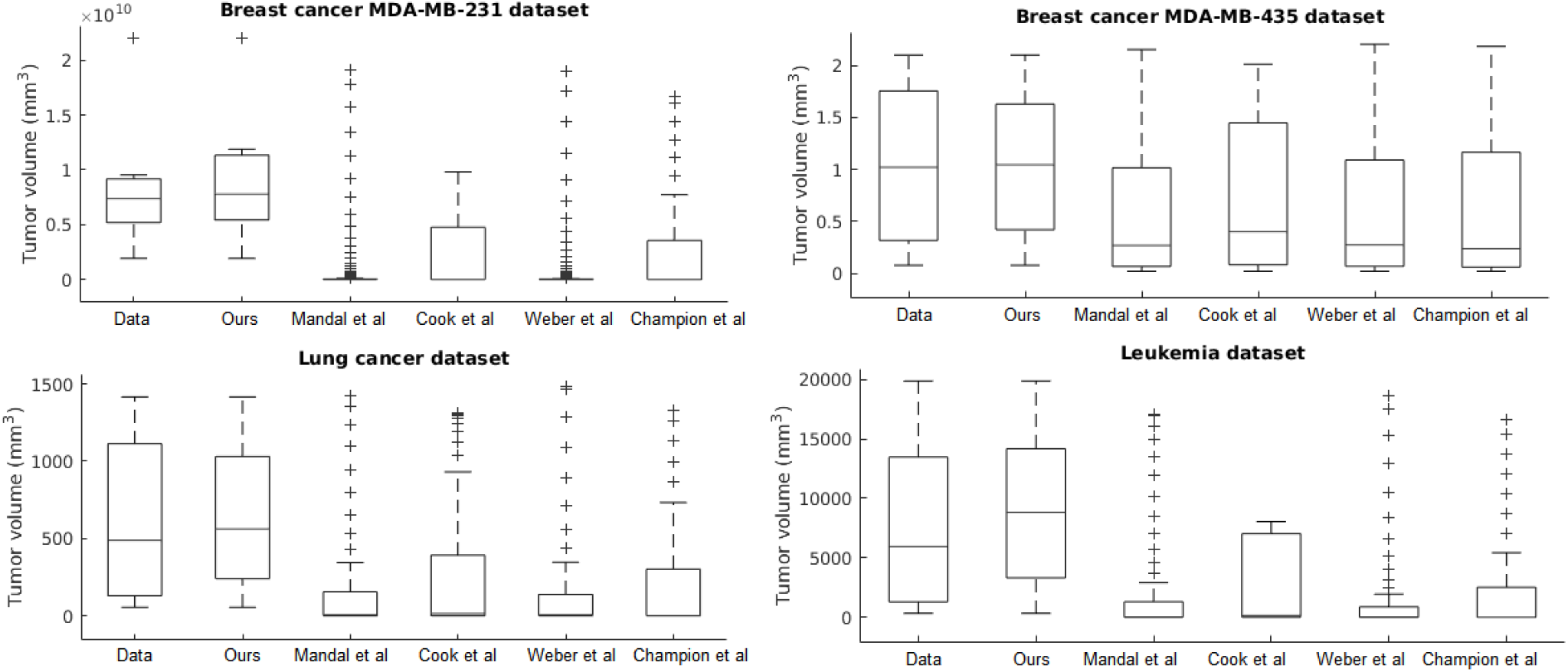
Evaluation of the data-driven relation learning systems on tumor growth: summary statistics.

#### 3.3.3 Extracting tumor phenotypic stage transitions

The next evaluation task looks at learning the mathematical relations describing the phenotypical transitions of tumors in breast cancer. For this experiment, we considered the study of 17 breast cancer patients in the study of Edgerton et al. (2011). Typically, in the breast cancer phenotypic state space, quiescent cancer cells (Q) can become proliferative (P) or apoptotic (A). Additionally non-necrotic cells become hypoxic if the oxygen supply drops below a threshold value. But, hypoxic cells can recover to their previous state or become necrotic, as shown in Macklin et al. (2012).

In this instantiation, we focus on a simplified 3-state phenotypic model (i.e. containing P, Q, A states). The transitions among tumor states are stochastic events generated by Poisson processes. Each of the data-driven relation learning systems is fed with timeseries of raw immunohistochemistry and morphometric data for each of the 17 tumor cases (see Edgerton et al. (2011), Tables S1 and S2) as following: cells cycle time *τ*_*P*_, cells apoptosis time *τ*_*A*_, proliferation index *PI* and apoptosis index *AI*. Given this timeseries input each system needs to infer the mathematical relations for *α*_*P*_, the mean Q - P transition rate, and *α*_*A*_, the Q - A transition rate, respectively (see Figure 6). Their analytical form the state transition is given by:

**Figure 6.**
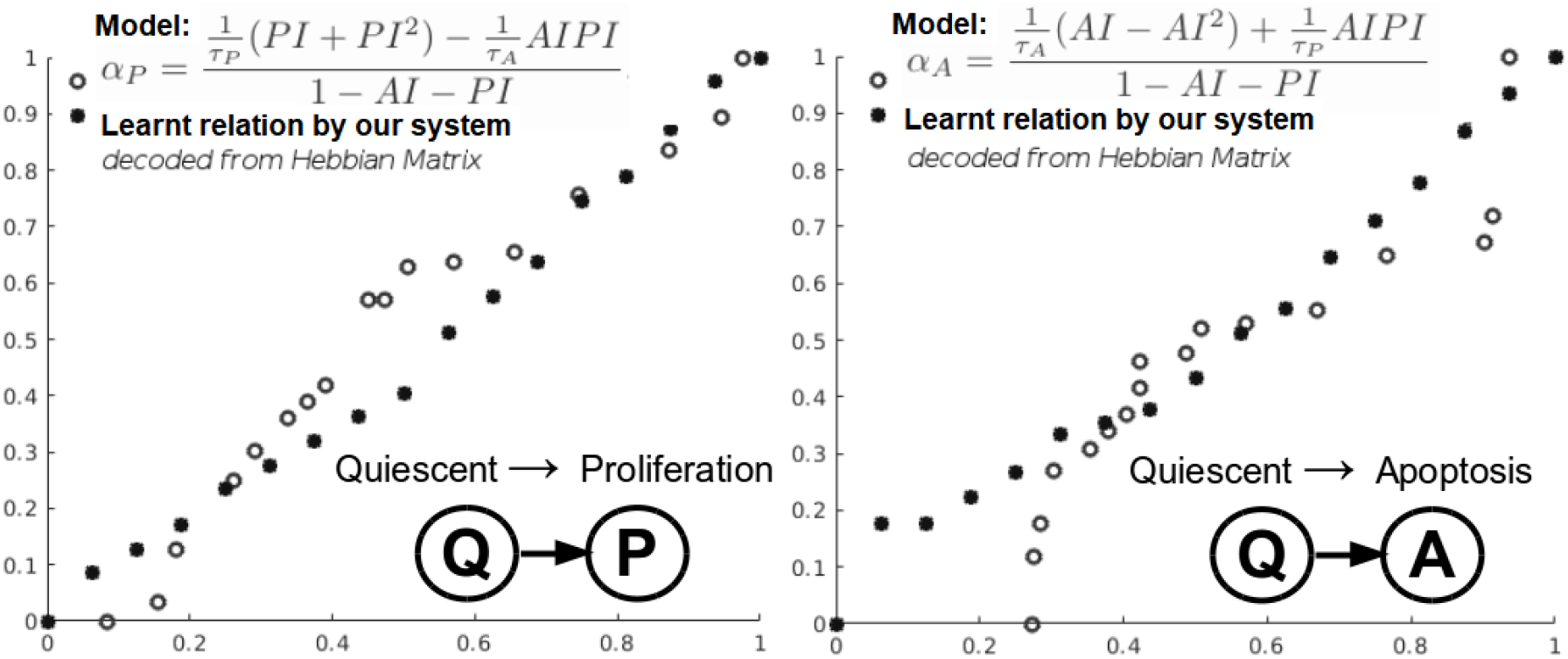
Learning cancer cells phenotypical states transitions mathematical relations.

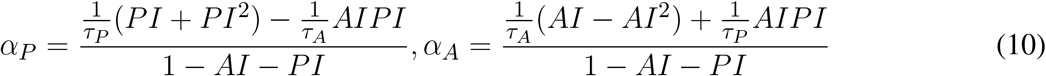

Quiescent (Q) to apoptosis (A) and quiescent (Q) to proliferation (P) state transitions of cancer cells are depicted in Figure 6, where we also present the relation that our system learnt. Both in Figure 6 and Table 5, we can see that our system is able to recover the correct underlying mathematical function with respect to ground truth (clinically extracted and modelled Equation 10 from the study of Macklin et al. (2012)).

**Table 5.** Evaluation of the data-driven relation learning systems for extracting phenotypic transitions relations.

|  | Evaluation Metrics |  |  |
| --- | --- | --- | --- |
| State Transition/System | SSE | RMSE | sMAPE |
| *Quiescent(Q) to Proliferation(P) Transition Relation* |  |  |  |
| Cook et al. | 0.820 | 0.240 | 0.190 |
| Weber et al. | 0.865 | 0.294 | 0.196 |
| Mandal et al. | 0.904 | 0.320 | 0.214 |
| Champion et al. | 0.845 | 0.274 | 0.189 |
| **Our system** | **0.750** | **0.210** | **0.172** |
| *Quiescent(Q) to Apoptosis(A) Transition Relation* |  |  |  |
| Cook et al. | 0.421 | 0.162 | 0.140 |
| Weber et al. | 0.484 | 0.178 | 0.154 |
| Mandal et al. | 0.490 | 0.182 | 0.151 |
| Champion et al. | 0.441 | 0.166 | 0.147 |
| **Our system** | **0.398** | **0.153** | **0.131** |

Note that none of the evaluated system had prior knowledge of the data distribution or biological assumptions. To have a more detailed overview on the capabilities of our system to capture phenotypic dynamics please refer to Axenie and Kurz (2020c).

#### 3.3.4 Simultaneously extracting drug-perturbed tumor growth and drug pharmacokinetics

Chemotherapy use in the neoadjuvant and adjuvant settings generally provides the same long-term outcome (de Wiel et al. (2017)). But what is the best choice for a particular patient? This question points at those quantifiable patient-specific factors (e.g. tumor growth curve under chemotherapy, drug pharmacokinetics) that influence the sequencing of chemotherapy and surgery in a therapy plan. A large variety of breast cancer tumor growth patterns used in cancer treatments planning were identified experimentally and clinically, and modelled over the years Gerlee (2013). Additionally, progress in pharmacokinetic modelling allowed clinicians to investigate the effect of covariates in drug administration, as shown in the work of Zaheed et al. (2019). Considering breast cancer, Paclitaxel is a typical drug choice with broad use in monothereapy as well as immune-combined therapies Stage et al. (2018).

In the current section, we present the experimental results of all the evaluated systems and consider: a) accuracy in learning the chemotherapy-perturbed tumor growth model, and b) accuracy in learning the pharmacokinetics of the chemotoxic drug (i.e. Paclitaxel) dose. For the tumor growth curve extraction, we considered four cell lines of breast cancer (i.e. MDA-MB-231, MDA-MB-435, MCF-7, LM2-LUC+ cell lines - see Table1). The evaluation results of the systems in the perturbed tumor growth scenario are provided in Figure 7.

**Figure 7.**
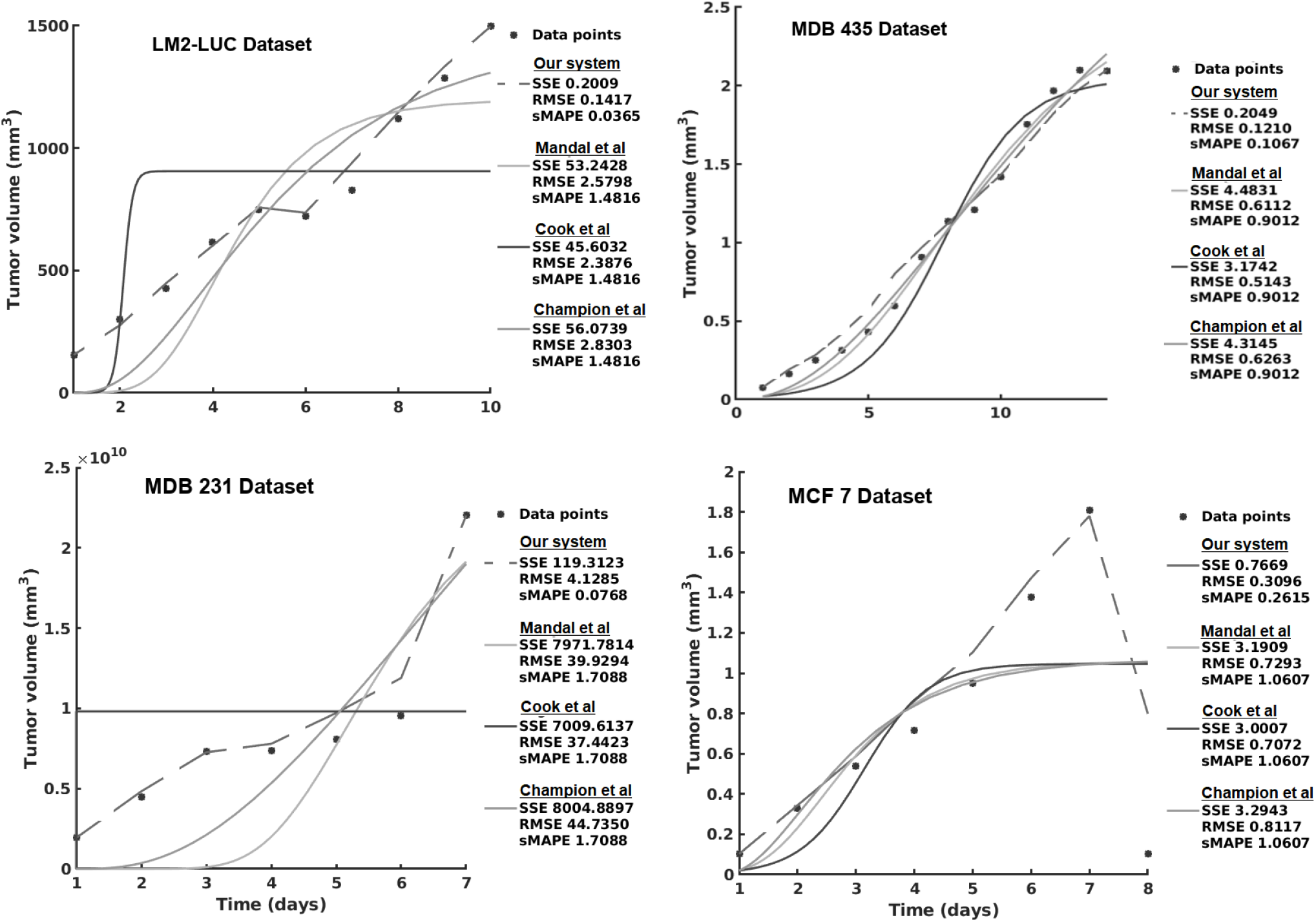
Evaluation of the data-driven relation learning system on perturbed tumor growth: accuracy evaluation. The decrease in the MCF7 Dataset is due to a high dose chemotherapy administration and demonstrates the adaptivity of the methods to cope such abnormal growth behaviours.

Table 6 presents the results using SVM and the different versions of DL. We can see that usually, vanilla DL outperforms SVM. DL is a more complex model, as well as uses more input data, so this result is expected. Once we add the augmentation from SVM, the model has a comparable performance to SVM. Our theory is that this is caused by DL learning to imitate SVM instead of real data. Once we add noise to the augmentation, the data becomes more realistic, and usually yields improvements in performance.

**Table 6.**
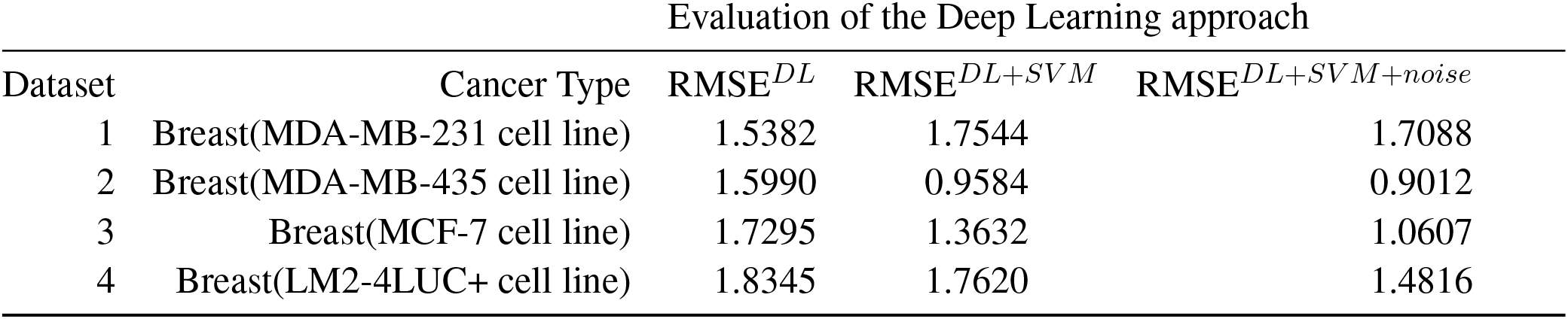
Description of the DL approach inspired by Champion et al. (2019).

| Evaluation of the Deep Learning approach |  |  |  |  |
| --- | --- | --- | --- | --- |
| Dataset | Cancer Type | RMSE<sup>DL</sup> | RMSE<sup>DL+SVM</sup> | RMSE<sup>DL+SVM+noise</sup> |
| 1 | Breast(MDA-MB-231 cell line) | 1.5382 | 1.7544 | 1.7088 |
| 2 | Breast(MDA-MB-435 cell line) | 1.5990 | 0.9584 | 0.9012 |
| 3 | Breast(MCF-7 cell line) | 1.7295 | 1.3632 | 1.0607 |
| 4 | Breast(LM2-4LUC+ cell line) | 1.8345 | 1.7620 | 1.4816 |

Note that for all of the evaluation datasets, the best performing DL approach (i.e. inspired by Champion et al. (2019)) is the combined DL - SVM - noise configuration.

For the pharmacokinetics learning experiments, we used the data from the computational model of intracellular pharmacokinetics of Paclitaxel of Kuh et al. (2000) describing the kinetics of Paclitaxel uptake, binding, and efflux from cancer cells in both intracellular and extracellular contexts.

As one can see in Figure 8 - left panel, the intracellular concentration kinetics of Paclitaxel is highly nonlinear. Our system is able to extract the underlying function describing the data without any assumption about the data and other prior information, opposite to the model from Kuh et al. (2000). Interestingly, our system captured a relevant effect consistent with multiple Paclitaxel studies Stage et al. (2018). Namely, that the intracellular concentration increased with time and approached plateau levels, with the longest time to reach plateau levels at the lowest extracellular concentration - as shown in Figure 8.

**Figure 8.**
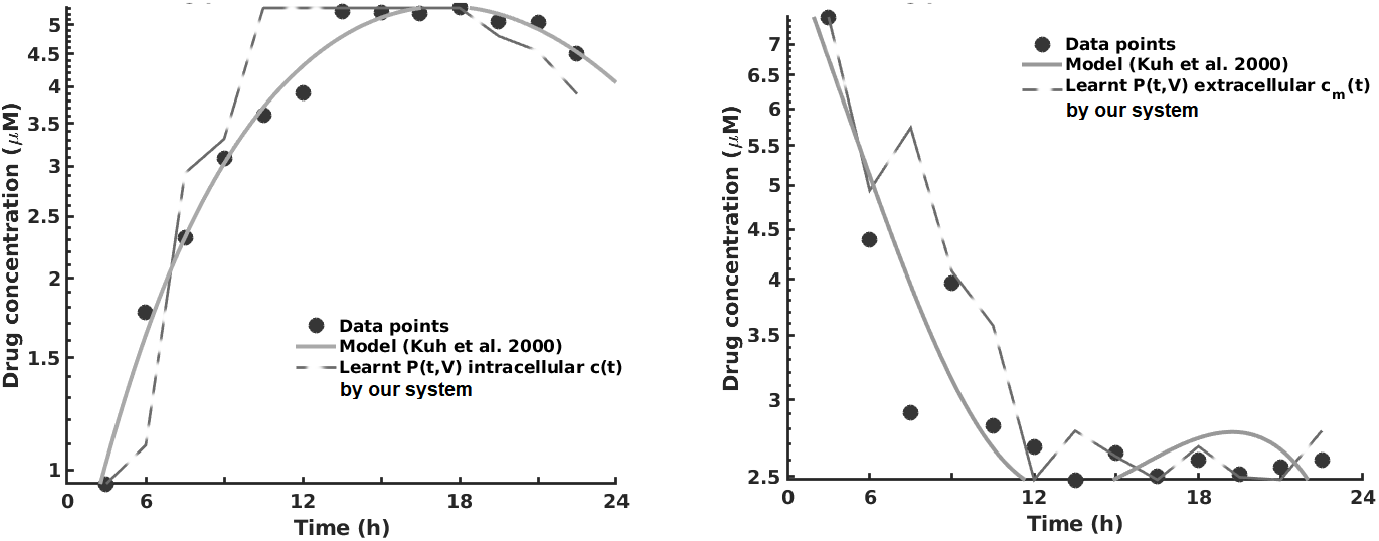
Learning the pharmacokinetics of the intracellular (left) and extracellular (right) Paclitaxel concentration. Data from Kuh et al. (2000), log scale plot.

Analysing the extracellular concentration in Figure 8 - right panel, we can see that our system extracted the trend and the individual variation of drug concentration after the administration of the drug (i.e. in the first 6h) and learnt an accurate fit without any priors or other biological assumptions. Interestingly, our system captured the fact that the intracellular drug concentration increased linearly with extracellular concentration decrease, as shown in Figure 8.

The overall evaluation of pharmacokinetics learning is given in Table 7.

**Table 7.** Evaluation of the data-driven relation learning systems for pharmacokinetics extraction.

| Pharmacokinetics data/System | Evaluation Metrics |  |  |
| --- | --- | --- | --- |
|  | SSE | RMSE | sMAPE |
| *Intracellular Paclitaxel* |  |  |  |
| Cook et al. | 0.6234 | 0.2812 | 0.1487 |
| Mandal et al. | 0.6743 | 0.3046 | 0.1602 |
| Champion et al. | 0.6539 | 0.2607 | 0.1500 |
| **Our system** | **0.5960** | **0.2141** | **0.1403** |
| *Extracellular Paclitaxel* |  |  |  |
| Cook et al. | 0.5676 | 0.2341 | 0.1213 |
| Mandal et al. | 0.6128 | 0.2974 | 0.1366 |
| Champion et al. | 0.5790 | 0.2633 | 0.1156 |
| **Our system** | **0.5484** | **0.2054** | **0.1068** |

In this series of experiments all of the systems learnt that changes in cell number were represented by changes in volume which: 1) increased with time at low initial total extracellular drug concentrations due to continued cell proliferation and 2) decreased with time at high initial total extracellular drug concentrations due to the anti-proliferative and/or cytotoxic drug effects, as reported in Kuh et al. (2000). In order to asses the impact the predictions have upon therapy sequencing (i.e. neoadjuvant vs. adjuvant chemotherapy) please refer to Axenie and Kurz (2020a).

#### 3.3.5 Predicting tumor growth/recession under chemotherapy

In the last series of experiments, we used real patient data from the I-SPY 1 TRIAL: ACRIN 6657 Yee et al. (2020). Data for the 136 patients treated for breast cancer in the IPSY-1 clinical trial was obtained from the cancer imaging archive ^6^ and the Breast Imaging Research Program at UCSF. The timeseries data contained only the largest tumor volume from MRI measured before therapy, 1 to 3 days after therapy, between therapy cycles, and before surgery, respectively. To summarize, the properties of the dataset are depicted in Figure 9.

**Figure 9.**
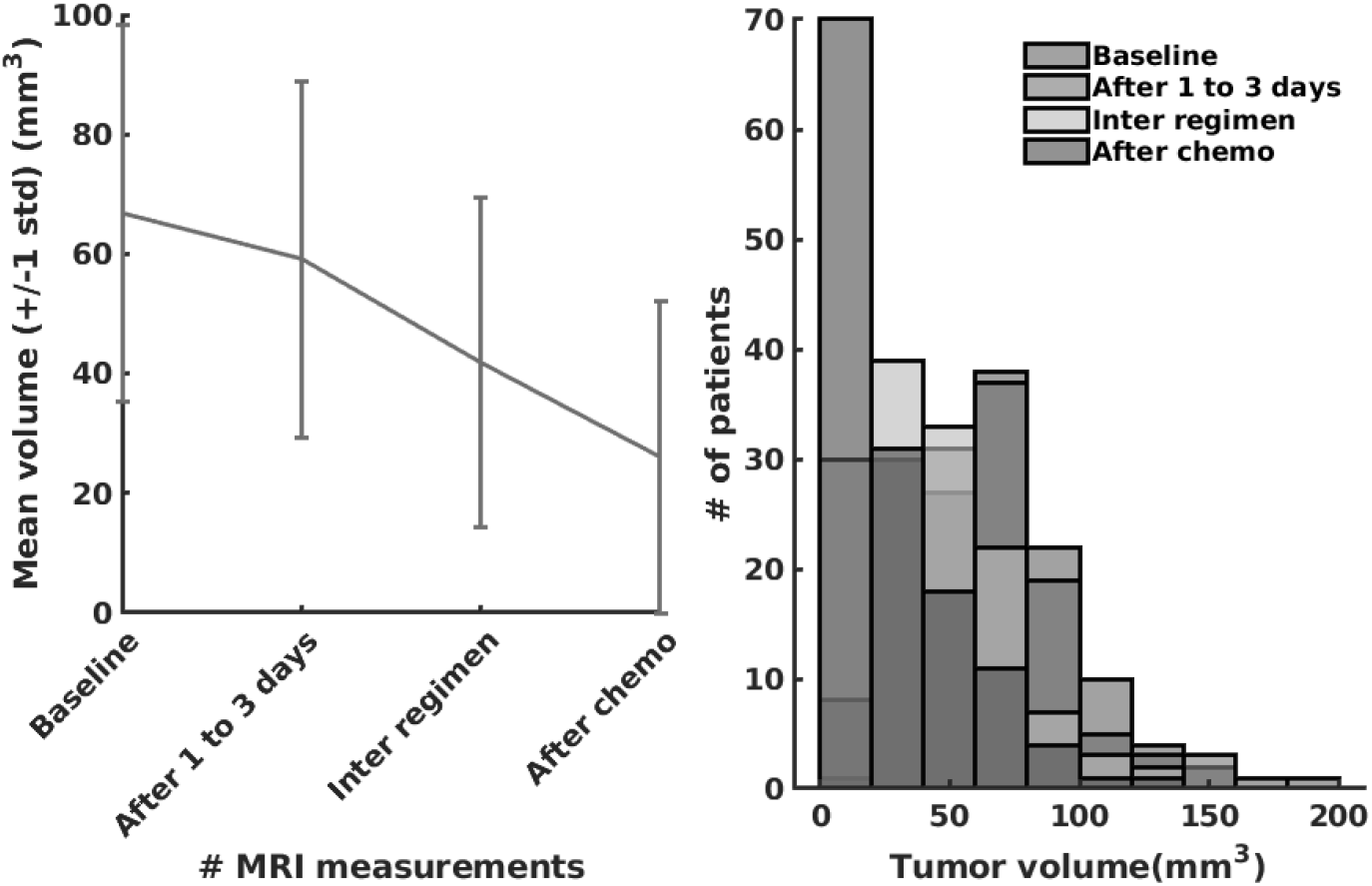
The I-SPY2 Trial Dataset properties.

As we can observe in Table 8, our system learns a superior fit to the tumor growth data, with respect to the other systems, despite the limited number of samples (i.e. 7 data points for MDA-MD-231 cell line dataset and up to 14 data points for MDA-MD-435 cell line dataset). Important, to note, that when analyzing tumor growth functions and response under chemotherapy, we faced the high variability among patients given by the typical constellation of hormone receptors indicators (i.e. HR and HER2neu, which covered the full spectrum of positive and negative values) for positive and negative prognosis. All data-driven learning systems capture such aspects to some extent. Our system learns a superior fit overall the three metrics, capturing the intrinsic impact chemotherapy has upon the tumor growth function, despite the limited number of samples (i.e. 4 data points of the dataset overall evaluation dataset of 20% of patients). An extended evaluation of our system on a broader set of datasets for therapy outcome prediction is given in Kurz and Axenie (2020).

**Table 8.** Evaluation of the data-driven relation learning systems on real patient breast cancer data.

| Dataset/Model | Evaluation Metrics |  |  |
| --- | --- | --- | --- |
|  | SSE | RMSE | sMAPE |
| *I-SPY2 Trial* Yee et al. (2020) |  |  |  |
| Cook et al. | 1.3735 | 1.1439 | 0.1133 |
| Mandal et al. | 2.963 | 1.0963 | 0.7834 |
| Champion et al. | 2.0747 | 1.04100 | 0.1073 |
| **Our system** | **0.8650** | **0.4650** | **0.0389** |

## 4 DISCUSSION

We complement the quantitative evaluation in the previous section with an analysis of the the most important features of all the systems capable to extract mathematical relations in the aforementioned clinical oncology tasks. As the performance evaluation was done in the previous section, we will now focus on other specific comparison terms relevant for the adoption of such systems in clinical practice.

One initial aspect is the design and functionality. Either using distributed representations Cook et al. (2010); Weber and Wermter (2007); Champion et al. (2019) or compact mathematical forms Mandal and Cichocki (2013), all methods encoded the input variables in a new representation to facilitate computation. At this level, employing neural network dynamics Cook et al. (2010); Weber and Wermter (2007) or pure mathematical multivariate optimisation Mandal and Cichocki (2013); Champion et al. (2019), the solution was obtained through iterative processes that converged to consistent representations of the data. Our system employs a light-weight learning mechanism, offering a transparent processing scheme and human-understandable representation of the learnt relations.

A second aspect refers to the amount of prior information embedded by the designer in the system. It is typical that, depending on the instantiation, a new set of parameters is needed, making the models less flexible. Although less intuitive, the pure mathematical approaches Mandal and Cichocki (2013) (i.e. using CCA) need less tuning effort due to the fact that their parameters are the result of an optimisation procedure. On the other side, the neural network approaches Cook et al. (2010); Weber and Wermter (2007); Champion et al. (2019) need a more judicious parameter tuning, as their dynamics are more sensitive, and can either reach instability (e.g. recurrent networks) or local minima. Except parametrisation, prior information about inputs is generally needed when instantiating the system for a certain scenario. Sensory values bounds and probability distributions must be explicitly encoded in the models through explicit distribution of the input space across neurons in Cook et al. (2010); Weber and Wermter (2007), linear coefficients in vector combinations Mandal and Cichocki (2013), or standardisation routines of input variables Champion et al. (2019). Our system exploits only the available data to simultaneously extract the data distribution and the underlying mathematical relation governing tumor growth processes.

A third aspect relevant to the analysis is the stability and robustness of the obtained representation. The representation of the hidden relation: 1) can be encoded in a weight matrix Cook et al. (2010); Weber and Wermter (2007) such that, after learning, given new input, the representation is continuously refined to accommodate new inputs; 2) can be fixed in vector directions of random variables requiring a new iterative algorithm run from initial conditions to accommodate new input Mandal and Cichocki (2013); or 3) can be obtained as an optimisation process given the new available input signals Champion et al. (2019). Given initial conditions, prior knowledge and an optimisation criteria Mandal and Cichocki (2013) or a recurrent relaxation process towards a point attractor Cook et al. (2010); Weber and Wermter (2007); Champion et al. (2019) are required to reach a desired tolerance. Our system exploits the temporal regularities among tumor growth data covariates, to learn the governing relations using a robust distributed representation of each data quantity.

The capability to handle noisy data, is an important aspect concerning the applicability in real-world scenarios. Using either computational mechanisms for de-noising Cook et al. (2010); Weber and Wermter (2007), iterative updates to minimise a distance metric Mandal and Cichocki (2013), or optimisation Champion et al. (2019), each method is capable to cope with moderate amounts of noise. Despite this, some methods have intrinsic methods to cope with noisy data intrinsically, through their dynamics, by recurrently propagating correct estimates and balancing new samples Cook et al. (2010). The distributed representation used in our system ensures that the system is robust to noise, and the local learning rules ensure fast convergence on real world data — as our experiments demonstrated.

Another relevant feature is the capability to infer (i.e. predict / anticipate) missing quantities once the mathematical relation is learned. The capability to use the learned relations to determine missing quantities is not available in all presented systems, such as the system of Mandal and Cichocki (2013). This is due to the fact that the divergence and correlation coefficient expressions might be non-invertible functions that support a simple pass through of available values to extract missing ones. On the other side, using either the learned co-activation weight matrix Cook et al. (2010); Weber and Wermter (2007), or the known standard deviations of the canonical variants Champion et al. (2019) some systems are able to predict missing quantities. Our system stores learnt mathematical relations in the Hebbian matrix which can be used bi-directionally to recover missing quantities on one side of the input given the other available quantity.

Finally, due to the fact that all methods re-encode the real-world values in new representation, it is important to study the capability to decode the learned representation and subsequently measure the precision of the learned representation. Although not explicitly treated in the presented systems, decoding the extracted representations is not trivial. Using a tiled mapping of the input values along the neural network representations, the system of Cook et al. (2010) decoded the encoded value in activity patterns by simply computing the distribution of the input space over the neural population units, while Weber and Wermter (2007) used a simple winner-take-all readout given that the representation was constrained to have a uniquely defined mapping. Given that the model learns the relations in data space through optimisation processes, as in the system of Champion et al. (2019), one can use learned curves to simply project available sensory values through the learned function to get the second value, as the scale is preserved. Albeit its capability to precisely extract nonlinear relations from high-dimensional random datasets the system of Mandal and Cichocki (2013) cannot provide any readout mechanisms to support a proper decoded representation of the extracted relations. This is due to the fact that the method cannot recover the sign and scale of the relations. The human-understandable relation learnt by our system is efficiently decoded from the Hebbian matrix back to real-world values. As our experiments demonstrate the approach introduced through our system excels in capturing the peculiarities clinical data carry.

## 5 CONCLUSION

Data-driven approaches to improve decision making in clinical oncology are now going beyond diagnosis. From early detection of infiltrating tumors to unperturbed tumor growth phenotypic staging, and from pharmacokinetics-dictated therapy planning to treatment outcome, data-driven tools capable of learning hidden correlations in the data are now taking the foreground in mathematical and computational oncology. Our study introduces a novel framework and versatile system capable of learning physical and mathematical relations in heterogeneous oncology data. Together with a lightweight and transparent computational substrate, our system provides human-understandable solutions. This is achieved by capturing the distribution of the data in order to achieve superior fit and prediction capabilities between and withing cancer types. Supported by an exhaustive evaluation on in-vitro and in-vivo data, against state-of-the-art machine learning and deep learning systems, the proposed system stands out as a promising candidate for clinical adoption. Mathematical and computational oncology is an emerging field where efficient, transparent, and understandable data-driven systems hold the promise of paving the way to individualized therapy. But this can only be achieved by capturing the peculiarities of a patient’s tumor across scales and data types.

## AUTHOR CONTRIBUTIONS

DK designed the research, collected clinical datasets, and performed data analysis of clinical studies used in the experiments. CSS developed the source code for the experiments and the analysis. CA designed the research and developed the source code for the experiments.

## Footnotes

1 A copy of the used datasets along with the study generating them is included in the codebase associated with the manuscript.

2 Experimental codebase: https://gitlab.com/akii-microlab/math-comp-ml

6 https://wiki.cancerimagingarchive.net/display/Public/ISPY1

